# A convergent depression vulnerability pathway encoded by emergent spatiotemporal dynamics

**DOI:** 10.1101/154708

**Authors:** Rainbo Hultman, Kyle Ulrich, Benjamin D. Sachs, Cameron Blount, David E. Carlson, Nkemdilim Ndubuizu, Rosemary C. Bagot, Eric Parise, Mai-Anh T. Vu, Joyce Wang, Alcino J. Silva, Karl Deisseroth, Stephen Mague, Marc G. Caron, Eric J. Nestler, Lawrence Carin, Kafui Dzirasa

**Affiliations:** Dept. of Psychiatry and Behavioral Sciences; Dept. of Neurobiology; Center for Neuroengineering; Dept. of Cell Biology; Dept. of Neurosurgery; Duke Institute for Brain Sciences, Duke University Medical Center, Durham, North Carolina 27710, USA; Dept. of Civil and Electrical Engineering; Dept. of Electrical and Computer Engineering; Dept. of Biomedical Engineering, Duke University, Durham North Carolina 22208, USA; Department of Psychological and Brain Sciences, Villanova University, Villanova, PA, 19085, USA.; Fishberg, Department of Neuroscience, Friedman Brain Institute, Ichan School of Medicine at Mount Sanai, 1 Gustave L. Levy Place, New York, New York 10029, USA.; Depts. of Neurobiology, Psychiatry & Behavioral Sciences and Psychology, Integrative Center for Learning and Memory, Brain Research Institute, University of California, Los Angeles, California 90095, USA.; Depts. of Bioengineering and Psychiatry and Howard Hughes Medical Institute, Stanford University, Stanford, California 94305, USA.

## Abstract

Fluctuations in brain local field potential (LFP) oscillations reflect emergent network-level signals that mediate behavior. Cracking the code whereby these LFP oscillations coordinate in time and space (spatiotemporal dynamics) to represent complex behaviors would provide fundamental insights into how the brain signals emotional processes at the network level. Here we use machine learning to integrate LFP activity acquired concurrently from seven cortical and subcortical brain regions into an analytical model that predicts the emergence of depression-related behavioral dysfunction across individual mice subjected to chronic social defeat stress. We uncover a spatiotemporal dynamic network in which activity originates in prefrontal cortex (PFC) and nucleus accumbens (NAc, ventral striatum), relays through amygdala and ventral tegmental area (VTA), and converges in ventral hippocampus (VHip). The activity of this network correlates with acute threat responses and brain-wide cellular firing, and it is enhanced in three independent molecular-, physiological-, and behavioral-based models of depression vulnerability. Finally, we use two antidepressant manipulations to demonstrate that this vulnerability network is biologically distinct from the networks that signal behavioral dysfunction after stress. Thus, corticostriatal to VHip-directed spatiotemporal dynamics organized at the network level are a novel convergent depression vulnerability pathway.

## Main Text

Major depressive disorder is the leading cause of disability in the world[1]. While stress contributes to the onset of depression[2, 3], only a fraction of individuals that experience stressful events develop behavioral pathology. Multiple factors including childhood trauma and alterations in several molecular pathways have been shown to increase disease risk[3, 4]; nevertheless, the neural pathways on which these factors converge to yield subthreshold changes that render individuals vulnerable to stress are unknown. Knowledge of these neural pathways would facilitate the development of novel diagnostic technologies that stratify disease risk as well as preventative therapeutics to reverse neural circuit endophenotypes that mediate vulnerability to depression. To achieve this aim it is essential to distinguish the neural alterations that confer vulnerability to depression from those that accompany the emergence of behavioral dysfunction.

Chronic social defeat stress (cSDS) is a widely validated pre-clinical model of depression[5-7]. In this paradigm, test mice are repeatedly exposed to larger aggressive CD1 mice. At the end of these exposures, test mice develop a depression-like behavioral state characterized by social avoidance, anhedonia- and anxiety-like behavior, and sleep/circadian dysregulation[6, 7]. Critically, only ㌬60% of C57 mice subjected to this paradigm exhibit susceptibility to developing this stress-induced syndrome. While the remaining ∼40% of mice subjected to cSDS exhibit resilience[6], susceptible and resilient mice experience the same degree of aggressive encounters. Thus, the cSDS paradigm provides a framework to probe putative basal network vulnerabilities that may exist in stress-vulnerable mice prior to stress exposure.

Multiple regions including subgenual cingulate cortex, amygdala, VHip, or VTA have been proposed to contribute to a putative depression brain network[5, 8-14]. Supporting this notion, functional magnetic resonance imaging (fMRI) studies in depressed subjects have discovered distinct functional connectivity alterations involving these brain regions that predict individual behavioral phenotypes and antidepressant treatment responses (i.e., pharmacology, psychotherapy, and transcranial magnetic stimulation)[15, 16]. However, our prior *in vivo* findings in genetic mouse models of depression and in mice exposed to cSDS suggest that depression-like behavioral dysfunction also arises at the level of circuit/network spatiotemporal dynamics, involving altered interactions of neural activity between spatially separated brain regions over time that are not captured by the fMRI timescale [14, 17, 18]. We postulated that a signature predicting depression vulnerability may exist at this dynamic circuit/network-level as well.

To test this hypothesis, we employed a transdisciplinary strategy integrating cSDS in mice, multi-circuit *in vivo* recordings from a subset of depression-related regions including prelimbic cortex (PrL_Cx), infralimbic cortex (IL_Cx), nucleus accumbens (NAc), central nucleus of the amygdala (CeA), basolateral amygdala (BLA), VTA, and VHip (Fig. 1a-b), a translational assay of neural circuit reactivity (Fig. 1b) [14, 19], and machine learning[20]. We selected this subset of brain regions since they have each been validated in contributing to depression-like behavior in multiple human and animal studies across several different research groups, and each region can be reliably targeted in mice using our multi-circuit recording technology[18, 21]. Our *in vivo* recording approach quantified both cellular activity and local field potentials (LFPs), which reflects the pooled activity of many neurons located up to 1mm from the electrode tip, their synaptic inputs, and their output signals[22].

**Figure 1:**
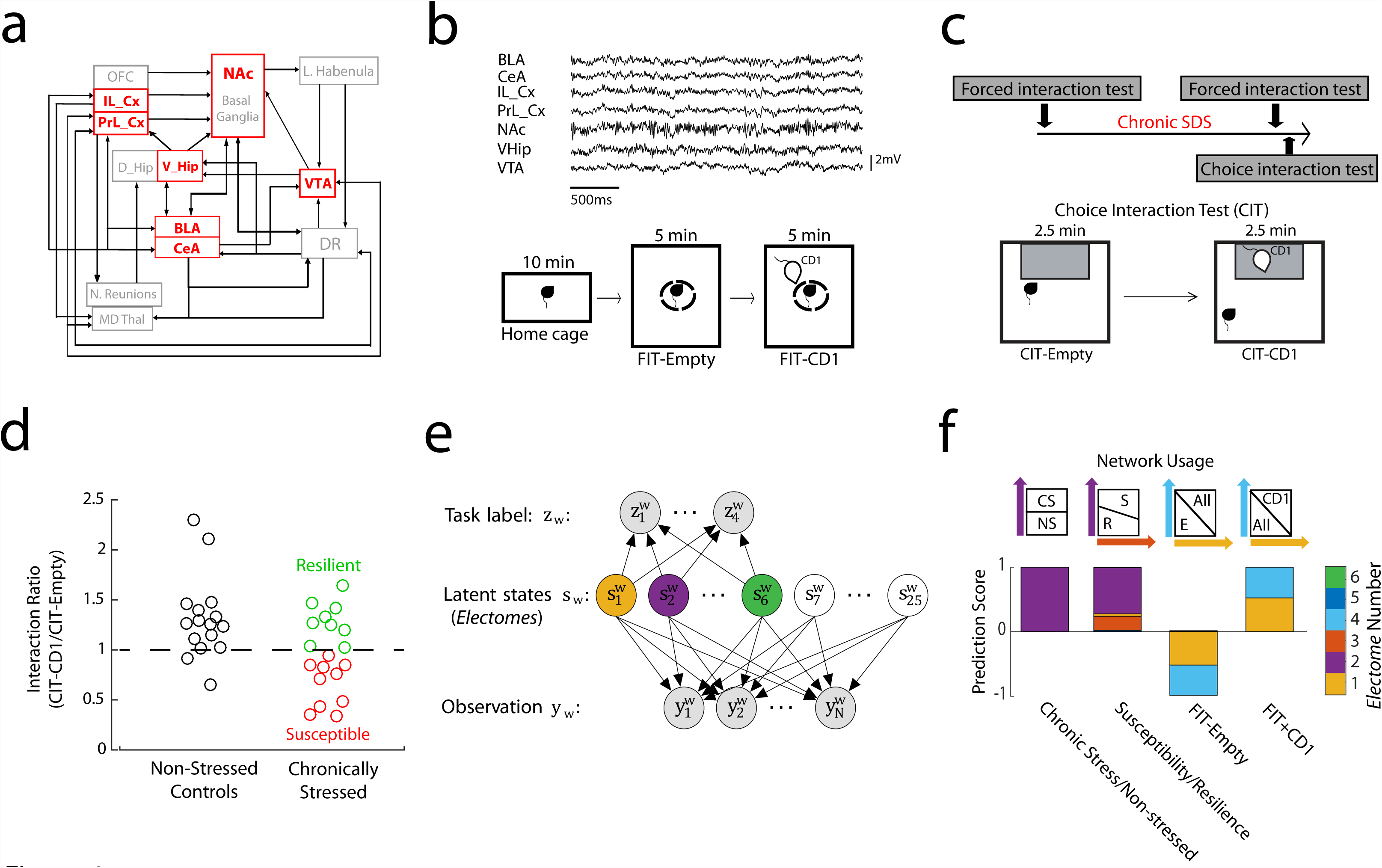
Identification of stress-related networks using machine learning. **a)** Partial wiring diagram describing structural interactions in mice across depression related brain regions. We performed multi-circuit recordings from the areas shown in red. **b)** Sample LFP traces recorded concurrently from seven brain areas: BLA, CeA, PrL_Cx, IL_Cx, NAc, VHip, and VTA (top). Homecage—Forced interaction test (FIT) used to probe brain network activity during: homecage, exposure to an empty cage while placed inside a small sub-chamber or to a cage containing a CD1 mouse while still inside small sub-chamber (bottom). **c)** Experimental timeline (top), and schematic of choice interaction test (CIT) classically used to identify susceptible vs. resilient mice after cSDS (bottom). **d)** Choice interaction ratios after 10 days of cSDS compared to non-stress controls. **e)** Cross-spectral factor analysis model where observations are brain features (LFP power, cross-area synchrony, and cross-area phase offsets) that are shared by latent states (networks). These networks coordinate distinct ‘emotional brain states’ represented by a given task label (i.e. susceptibility vs. resilience). We trained 25 descriptive latent networks. Six of these networks were also trained to be predictive **f)** Four networks/*Electomes* identified using a support vector machine jointly discriminated the stress states (Networks 1, 2, 3, and 4). Example support vectors are shown above.

We uncovered network-level spatiotemporal dynamic signatures that distinguish the neural alterations that confer vulnerability to depression from those that accompany the emergence of behavioral dysfunction after stress. We then utilized three independent mouse models of depression vulnerability to verify that one spatiotemporal dynamic network represents a convergent network-level vulnerability pathway for depression-related abnormalities. Finally, we used two distinct antidepressant manipulations to verify that this network underlying depression vulnerability is biologically distinct from the neural networks underlying the expression of depression-related behavioral dysfunction after stress exposure.

### Neural model of brain network function

To study the relationship between widespread spatiotemporal dynamics and depression pathology, we developed a novel probabilistic machine learning approach using LFP activity data recorded from seven brain regions across multiple frequencies. We term this novel approach “cross-spectral factor analysis” or CSFA (see Fig. 1e and supplemental methods for full details of CSFA model). Our CSFA approach yields a descriptive model, such that it discovers LFP patterns across regions that change together over seconds of time. The model is also predictive such that it discriminates the LFP patterns that are specific for a number of pre-specified behavioral variables. Our CSFA model parallels classic fMRI models that describe functional connectivity quantified by ultra-slow correlated activity across many brain regions over seconds of time. However, in contrast to fMRI models, our approach also enables the analysis of fast oscillatory electrical signals at the millisecond timescale. Indeed, the faster timescale features that contribute to the LFP patterns we observe include spectral power (LFP amplitude across frequencies), synchrony (a neural correlate of brain circuit function that quantifies how two LFPs correlate over a millisecond timescale), and phase-directionality (a neural correlate of information transfer which quantifies which of two synchronous LFPs leads the other), across many brain regions (see supplemental Fig. S1). We therefore refer to these LFP patterns as “*Electomes”,* “electrical functional connectomes”. Importantly, our CSFA model also yields an *Electome* activity score which indicates the activity of each *Electome* during each five second window of LFP data. A given brain area or circuit can belong to multiple *Electomes*, providing the opportunity for distinct *Electomes* to functionally interact to yield a global brain state.

### Neural networks signal depression vulnerability

Brain activity was recorded while animals were in their home cage and during a forced interaction test with an aggressive mouse (Fig. 1b). A subset of the mice were subjected to cSDS, and the post-stress susceptibility of these mice was characterized using the choice interaction test (Fig. 1c) which has been shown to reliably track the expression of the full depression-like behavioral syndrome[6]. All of the mice were then subjected to another home cage–forced interaction test recording. We trained our CSFA model using machine learning to determine the oscillatory signals that are modulated across time and discriminate: 1) all mice subjected to cSDS from non-stressed controls (post-cSDS; N=19 and 16 mice, respectively), 2) susceptible from resilient mice (post-cSDS; N=10 and 9 mice, respectively) and 3) activity recorded during different segments of the homecage--forced interaction test recordings (Fig. 1b, pre-cSDS and post-cSDS; N=44 total mice)[14, 19].

To determine which *Electomes* derived by our CSFA model discriminated these stress conditions, we used a support vector machine. We found that 4 out of the 25 specified *Electomes* discriminated these various behavioral conditions (Fig. 1f, see supplemental materials for detailed CSFA model description). For discriminating cSDS exposure, *Electome 2* activity was higher in mice subjected to cSDS than in non-stressed controls (Fig. 1f, see also Fig. 2d). *Electome* 2 was also higher in stress-susceptible mice compared to the resilient animals (Fig. 1f and 2d). This *Electome* was defined by co-modulated delta and beta oscillations, and oscillations in this network exhibited directionality largely from NAc to VHip and VTA (Fig. 2a-c). *Electome 3* was also higher in stress-susceptible mice compared to the resilient animals (Fig. 1f). This *Electome* was defined by co-modulated delta oscillations that exhibited directionality from PFC and NAc to BLA (Fig. 2a-c). Finally, *Electome 1* activity was enhanced by acute exposure to the CD1 mouse during the forced interaction test both before and after cSDS (Figs. 1f and 2d). *Electome 1* was largely defined by 8-20Hz oscillations that exhibit directionality from PFC and NAc to VHip (Fig. 2a-c). *Electome 4*, defined by local delta (1-4Hz) oscillations in VHip, did not show dramatic changes with behavioral conditions (Fig. 2d), although was nonetheless associated significantly with the forced interaction test (Fig. 1f).

**Figure 2:**
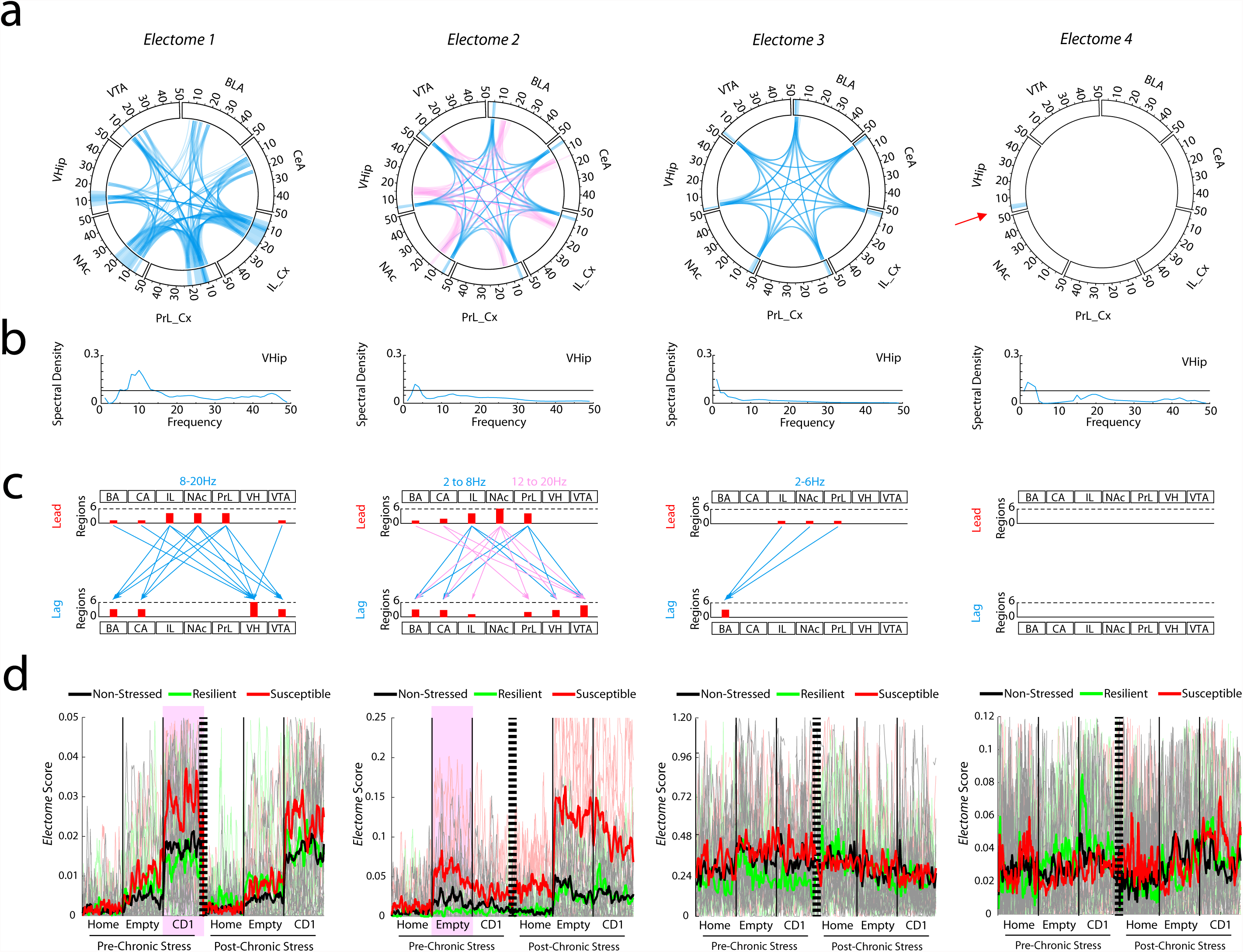
Four *Electome* networks signal distinct stress states. **a)** Power and coherence measures that compose each network. Brain areas and oscillatory frequency bands ranging from 1 to 50Hz are shown around the rim of the circle plot. Spectral power measures that contribute to each *Electome* are depicted by the highlights around the rim, and cross spectral (i.e., synchrony) measures are depicted by the lines connecting the brain regions through the center of the circle (*Electome* activity is shown at a relative spectral density threshold of 0.08). **b)** Example VHip spectral density plot for is shown for each *Electome*. **c)** Phase offset measures that define directionality within each *Electome* (phase activity is shown at a threshold of 0.1 radians). Histograms quantify the number of lead and lagging circuit interactions for each brain region. **d)** *Electome* activation during pre- and post-stress home cage and forced interaction test (FIT) recordings. The thick colored lines show the average across animals, while the thin lines in the background show the values from individual mice. Two *Electomes* (highlighted by purple) showed test-related statistical differences between susceptible and resilient mice prior to cSDS exposure (P<0.01; N=5-7 mice/group).

Strikingly, two of the *Electomes* signaled vulnerability prior to the cSDS experience. During acute exposure to the CD1 mouse (i.e. the second half of the forced interaction test), *Electome 1* was higher in stress-naïve mice that later exhibited susceptibility to cSDS than those mice that later exhibited resilience (P = 0.0057 for comparison of pre-stress *Electome 1* activity during the forced interaction with the CD1; Receiver Operating Characteristic AUC=0.86; N=9-10 mice per group; Fig. 2d). In contrast, *Electome 2* was higher in stress-naïve mice that later exhibited susceptibility, specifically when they were placed in the interaction chamber during the first half of the forced interaction test (P=9.7×10^−4^ for comparison of pre-stress *Electome 2* activity during forced interaction test-Empty; Receiver Operating Characteristic AUC=0.92; N=9-10 mice per group; Fig. 2d). We did not observe significant differences between stress-naive susceptible mice and stress-naïve resilient mice when they were in their home cage, or across any of the other *Electomes* (P>0.05 for all comparisons). Thus, *Electomes 1 and 2* were putative biomarkers of vulnerability since they distinguished the stress-naïve test mice that would later show behavioral dysfunction after cSDS from the mice that would later exhibit resilience. *Electome 2* was also a biomarker of the emergence of depression-related behavioral dysfunction as activity in this network was increased in the stress-susceptible mice compared to the resilient mice and the non-stress controls.

### *Electome* activity correlates with unit firing

Having identified these putative stress-related signatures, we set out to verify that the *Electomes* were a *bona fide* representation of biological activity and not simply abstract mathematical constructs. To do this, we tested whether *Electome* activity demonstrated a relationship to the activity of neurons recorded simultaneously from the seven brain regions, which is a clear reflection of biological function. Specifically, we quantified the activity of each of the 644 recorded neurons in 5 second bins and compared this activity to the activity of each *Electome* (Fig. 3a). To verify that the degree of correlation of the *Electomes* with cellular firing rates was meaningful and not due to random chance, we compared these results to randomly shuffled firing rates (see Supplemental Methods). Each of the four *Electomes* exhibited activity that correlated with 10-15% of the recorded cells (N=644 units pooled from all brain areas, Fig. 3b). Many neurons (250/644, 39%) showed activity that correlated with more than one *Electome*, and 21-51% of the neurons from each individual brain area correlated with at least one of the four *Electomes* (Fig. 3b). These data confirmed that the *Electomes* reflect network-level neural processes that emerge from cellular firing across large spatiotemporal distributions[23].

**Figure 3:**
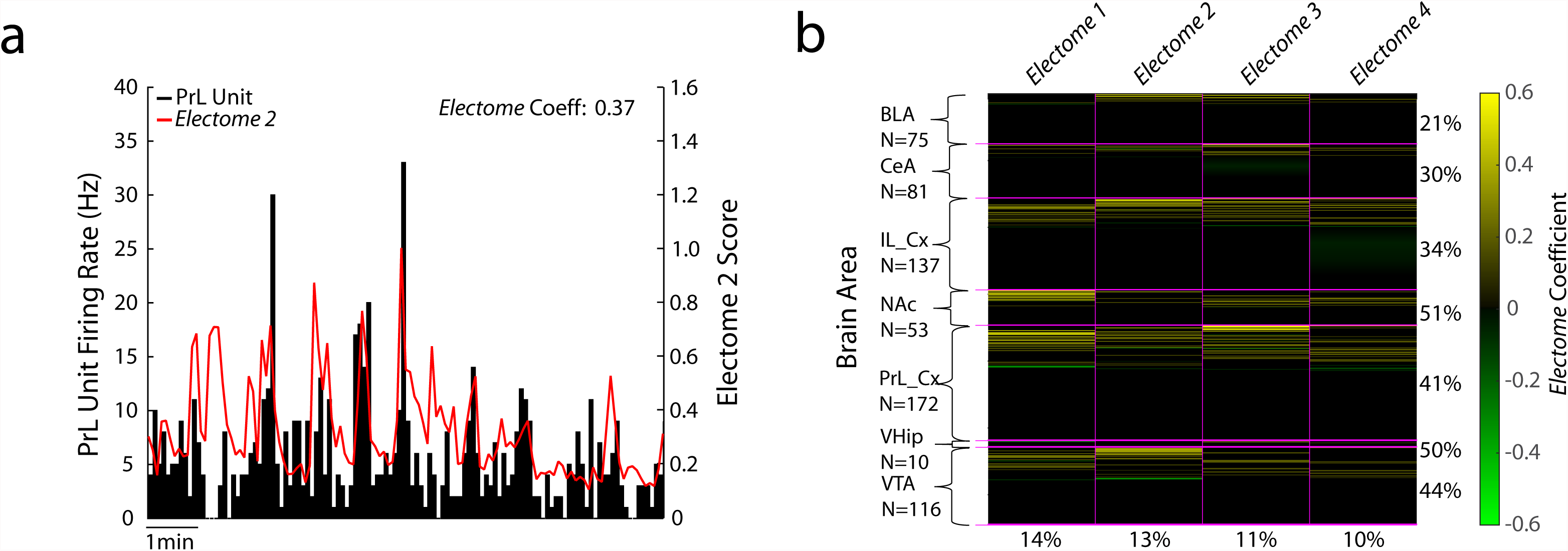
*Electome* network activity correlates with brain-wide cellular firing. **a)** Example of PFC neuron that showed significant firing relative to *Electome 2* activity in the home cage. **b)** Population firing relative to *Electome* activity (N = 644 cells). Yellow bars highlight units that showed firing that correlated with *Electome* activity. Green bars highlight units that showed anti-correlated firing relative to *Electome* activity.

### Enhanced *Electome* activity in a three independent models of vulnerability

We tested whether *Electome 1* and/or *Electome 2* indeed reflect a stress-vulnerability pathway that predicts susceptibility to future stress. We reasoned that, if these electrical patterns were truly reflective of general depression vulnerability mechanisms, then manipulations across many different levels of analysis implicated in depression vulnerability should also generate these electrical signatures. Thus, we subjected mice to overexpression of a susceptibility hub gene, chronic interferon-alpha treatment, or an early life stress, and directly tested whether these manipulations increased *Electome 1* or *Electome 2* activity.

We first exploited a molecular approach to enhance vulnerability in the cSDS model and then quantified the impact of this manipulation on the *Electomes*. Both *Electome* 1 and *Electome* 2 exhibited directionality towards VHip. Since we recently found that the *Sdk1* gene, which encodes the cell adhesion protein, sidekick 1, plays a central hub role in mediating susceptibility in VHip[24], we verified that Sdk1 overexpression in VHip increases stress vulnerability (P=0.0037; N=13-17 mice per group; Fig. 4a), as demonstrated previously. We then tested whether VHip-Sdk1 overexpression influences *Electome 1* or *Electome 2* activity, using a within subject design (Fig.4b). After an initial homecage—forced interaction test recording session, animals were injected intra-VHip with HSV-Sdk1-GFP or an HSV-GFP control vector (Fig. 4c). Two-days later, we repeated our neurophysiological recording protocol. By applying these recording data to the *Electome* model coefficients learned from our initial model in cSDS mice (Fig. 4d), we recovered *Electome* activity measures for the new testing sessions. Strikingly, VHip-Sdk1 overexpression, in the absence of stress, increased *Electome 1* activity during exposure to a CD1 mouse (P=0.037; N=5-7 mice; Fig. 4e). VHip-Sdk1 overexpression in the absence of chronic stress had no impact on *Electome 2*, nor did it yield the behavioral dysfunction that defines stress-susceptibility as observed previously (F_1,17_=1.03, P=0.32 for overexpression effect on social interaction; t_1,15_=0.07, P=0.95 for immobility time; see Fig. 4f-g)[24]. Thus, this molecular manipulation selectively induced the *Electome 1* spatiotemporal dynamic network and stress-vulnerability behavioral state that our computational model linked previously to enhanced vulnerability to cSDS in stress-naïve, wildtype mice.

**Figure 4:**
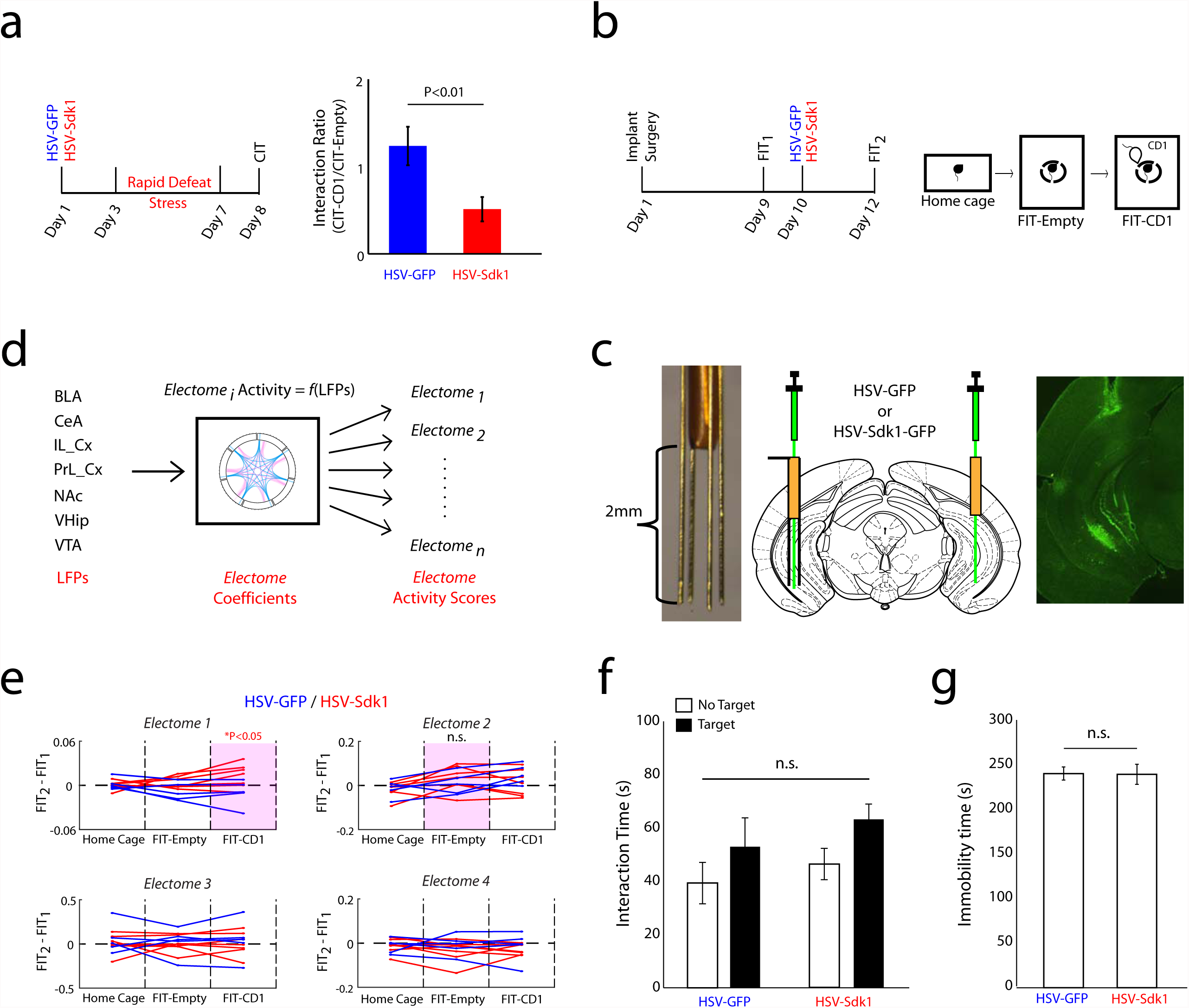
VHip-Sdk1 overexpression selectively increases *Electome 1* activity in stress-naïve mice. **a)** Mice were subjected social defeat twice daily for 4 days (i.e. accelerated defeat). The Sdk1 overexpression group exhibited increased susceptibility. **b)** Experimental schematic for neurophysiological recordings. **c)** Cannutrode enables site-specific viral injection in chronically implanted mice (left), surgical schematic (middle), and image showing GFP expression in chronically implanted mouse. **d)** LFPs recorded during the forced interaction test (FIT) were transformed using the initial CSFA *Electome* model/coefficients. **e)** Sdk1 overexpression in VHip increased *Electome 1* activity during the forced interaction test-CD1 (p<0.05 for comparison activity in HSV-Sdk1 and HSV-GFP mice using a one-tailed Wilcoxon rank-sum test). Purple boxes highlight network biomarkers of vulnerability to chronic stress in normal mice (see Fig. 2). **f-g)** Sdk1 overexpression had no significant effect on f) social interaction or g) immobility during a forced swim test in non-stressed mice.

Secondly, we tested whether a physiological manipulation, administration of interferon alpha (IFNα), a drug that induces a depression-like phenotype in humans[25], is sufficient to induce stress-related *Electome* activity (Fig. 5a). Prior studies have shown that chronic IFNα treatment induces a depressive-like behavioral syndrome in mice[26, 27]. Consistent with prior findings, we found that chronic IFNα treatment modestly reduced normal social behavior in a three-chamber social interaction test (F_1,18_=7.14, P=0.008; t_1,18_=2.1; P=0.03 for post-hoc testing of social time; N=10 mice per group; Fig. 5b). We also found that chronic IFNα treatment suppressed sucrose preference, verifying that this manipulation induced depression-related behavioral changes (F_1,16_=4.45, P=0.025; t_1,9_=2.3, P=0.024 for post-hoc testing of sucrose effect in Veh treated mice, N=10; t_1,7_=0.5, P=0.31 for sucrose effect in IFNα treated mice, N=8; Fig. 5d; see also supplemental Fig. S2).

**Figure 5:**
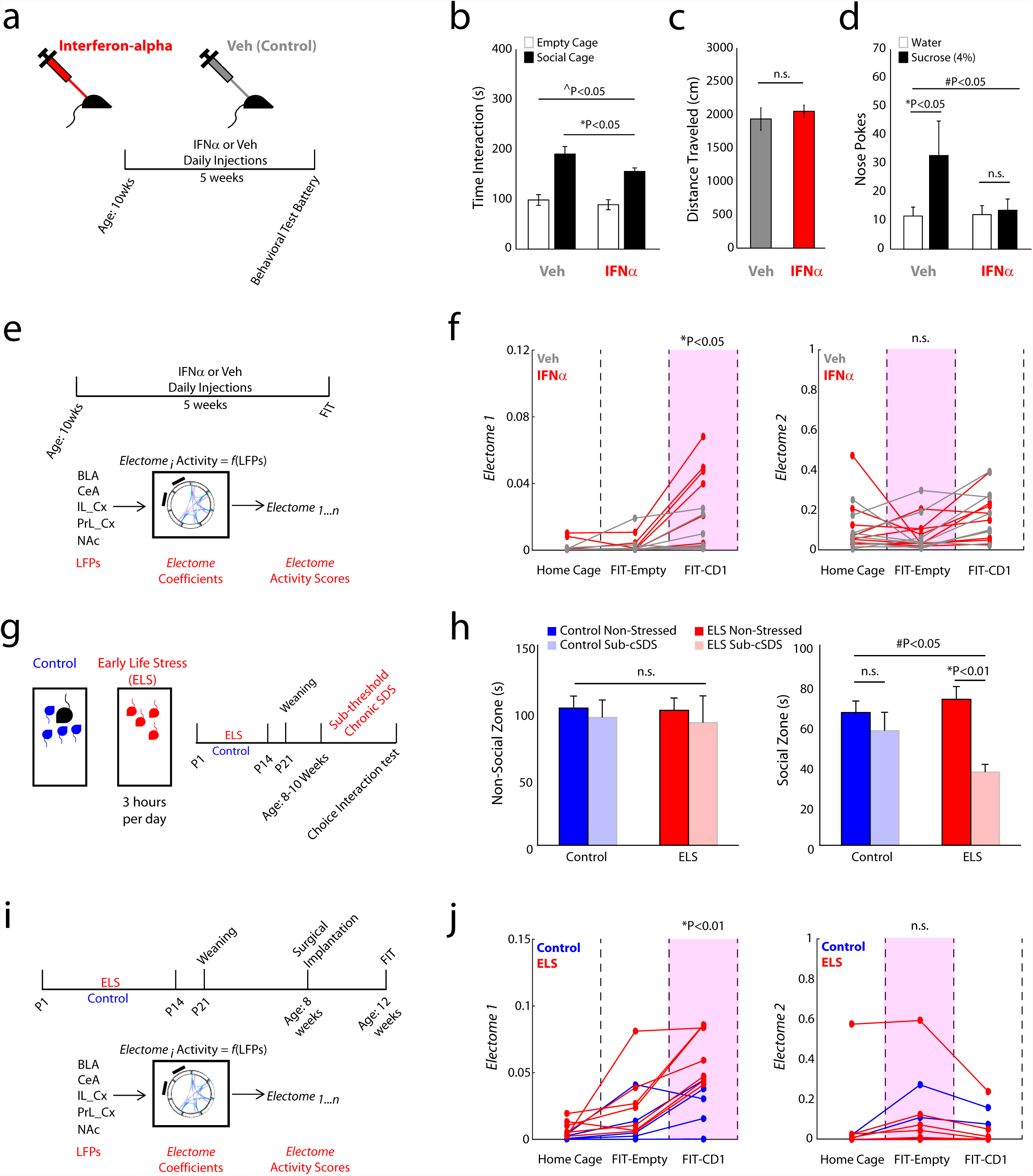
Enhanced *Electome 1* activity in two translational models of depression vulnerability. **a)** Experimental schematic. **b)** Chronic IFNα administration reduced social behavior in the classic three-chamber test (^P<0.05 for novel-mouse effect using two-way ANOVA, *P<0.05 using unpaired one-tailed t-test). **c)** No locomotor differences were observed in the open field (t_1,18_=0.599, P=0.56 using an unpaired two-tailed t-test; N=10 mice per group). **d)** Social behavior in three chamber social interaction test (#P<0.05 for sucrose effect using two-way ANOVA, *P<0.05 using paired one-tailed t-test). **e)** Schematic for neurophysiological experiments. **f)** Chronic IFNα treatment recapitulated the neurophysiological signature of stress vulnerability identified in *Electome 1*, but not *Electome 2*. **g)** Schematic for ELS paradigm and experimental timeline for neurophysiological testing**. h)** Impact of ELS and cSDS on social behavior (#P<0.05 for ELS x sub-threshold cSDS interaction effect using two-tailed two-way ANOVA; *P<0.05 using unpaired two-tailed t-test). **i)** Experimental schematic for *in vivo* recording experiments. **j)** ELS mice exhibited higher *Electome 1* activity during exposure to a CD1 mouse compared to normally reared controls. No difference was observed in *Electome 2* activity.

We then implanted C57 animals with micro-wire electrodes and treated with IFNα or vehicle. After 5 weeks of treatment, mice were subjected to the forced interaction test (Fig. 5e). Chronic IFNα treatment significantly increased *Electome 1* activity during CD1 exposure in the forced interaction test (P=0.041; N=8 mice per group; Fig. 5f). No difference was observed in *Electome 2* activity (P=0.323; Fig. 5f). Thus, IFNα treatment recapitulated the *Electome 1* spatiotemporal dynamic network we identified in the cSDS and Sdk1 models of depression vulnerability (see also supplemental Fig. S3). Notably, a powerful feature of our CSFA model is that once the original model and coefficients are learned, the same output features (*Electome* activity) can be determined from new data with LFP activity from only a subset of brain areas (Fig 5e). Thus, these mice were only implanted in the most technically accessible brain areas (PrL_Cx, IL_Cx, NAc, BLA, and CeA).

Thirdly, we sought to determine whether naturally occurring behavioral experiences that increase stress vulnerability also enhance our putative vulnerability network. Childhood trauma is a major risk factor for developing depression in adulthood[4]. Since maternal separation stress has been widely utilized as a mouse model of early life stress (ELS)[28, 29] (Fig. 5g), we first tested whether maternal separation stress was sufficient to render animals more vulnerable to stress in adulthood. ELS mice and their normally reared controls were subjected to a sub-threshold cSDS protocol where the animals were housed independently from the CD1 mice after each defeat. Mice subjected to ELS and sub-threshold cSDS exhibited the social avoidance that defines the stress-susceptible cSDS phenotype (F_1,34_=4.23, P=0.048; t_13_=5.43; P=0.0001 for post-hoc comparison of ELS/cSDS and ELS/non-stressed mice; N=7-8 per group; Fig 5h); However, neither ELS nor the sub-threshold cSDS exposure were sufficient to induce social avoidance on their own (t_15_=0.81; P=0.43 and t_18_=0.88; P=0.38, for comparison of normally reared/non-stressed mice to maternally-separated/non-stressed and normally reared/non-stressed, respectively, using FDR-corrected t-test; N=10 per group; Fig. 5h). No differences were observed in the interaction time with the non-social stimulus (F_1,34_=0.01, P=0.93 for ELS x sub-threshold cSDS interaction effect using two-tailed two-way ANOVA). Together, these findings verified that ELS increased vulnerability to adult stress.

We then implanted a new cohort of adult ELS mice and normally-reared controls with recording electrodes. After recovery, mice were subjected to the forced interaction test assay (Fig. 5i). By transforming the recorded LFP activity using our initial cSDS CSFA model and coefficients, we found that ELS increased *Electome 1* activity during exposure to the CD1 mouse (P=0.005 using one-tailed Wilcoxon Rank-sum test; N = 5-7 per group), with no effect on *Electome 2* activity (P=0.318 using one-tailed Wilcoxon Rank-sum test; Fig. 5j). Thus, ELS was sufficient to induce the vulnerability network signature in adult animals. Together, these findings confirmed that three independent molecular, physiological and behavioral manipulations that increase depression vulnerability in adult animals all converged on the same *Electome 1* network.

### The vulnerability network is distinct from depression-like behavior networks

Our initial cSDS CSFA model found that the network that signals latent stress *vulnerability* (*Electome 1*, prior to stress) was computationally distinct from the putative networks that signal the emergence of the depression-like behavior state in susceptible mice after cSDS (*Electome* 2 and *Electome* 3, post-stress). After validating *Electome 1* as a biological marker of depression vulnerability, we tested whether depression vulnerability was truly biologically distinct from depression-related behavioral abnormalities. We reasoned that if the *Electome 1* vulnerability signature was indeed mechanistically distinct from the networks underlying the pathological behavior state, biological manipulations that reverse depression-related behavioral abnormalities would fail to suppress *Electome 1* activity during our forced interaction test assay. Thus, we selected two distinct manipulations that have been shown to exert antidepressant effects in both humans and rodent models.

Deep brain stimulation (DBS) of subgenual cingulate cortex (Brodmann area 25, BA25) induces antidepressant effects in select clinical populations with depression[10, 30]. Critically, direct stimulation of left IL_Cx (the rodent homologue of BA25) exhibits antidepressant-like effects in the cSDS model as well[31]. To test the impact of left IL_Cx stimulation on *Electome* activity, we infected animals with a stabilized step-function opsin (SSFO, AAV-CaMKII-SSFO) in IL_Cx (Fig. 6a). When stimulated by blue light, SSFO induces increased firing of neurons for over 20 minutes[32]. Control animals were infected with a sham virus (AAV-Ef1a-DIO-SSFO), which does not express the opsin (Fig. 6a-b). We then implanted animals with electrodes and recorded their LFP activity in the forced interaction test immediately after blue light stimulation (Fig. 6b, see also supplemental Fig. S4). IL_Cx-DBS stimulation had no impact on *Electome 1* activity during exposure to the CD1 mouse (P=0.47 using rank-sum test; N=5-8 mice/group). However, IL_Cx-DBS stimulation did suppress *Electome 2* activity, even though the mice were stress-naïve (Fig. 6d, left; F_1,22_=6.3, P= 0.015). *Electome 3* activity was not suppressed by this manipulation (Fig. 6d, right; F_1,22_=0.99, P=0.17). Thus, as anticipated, IL_Cx-DBS stimulation had no impact on our depression-vulnerability signature.

**Figure 6:**
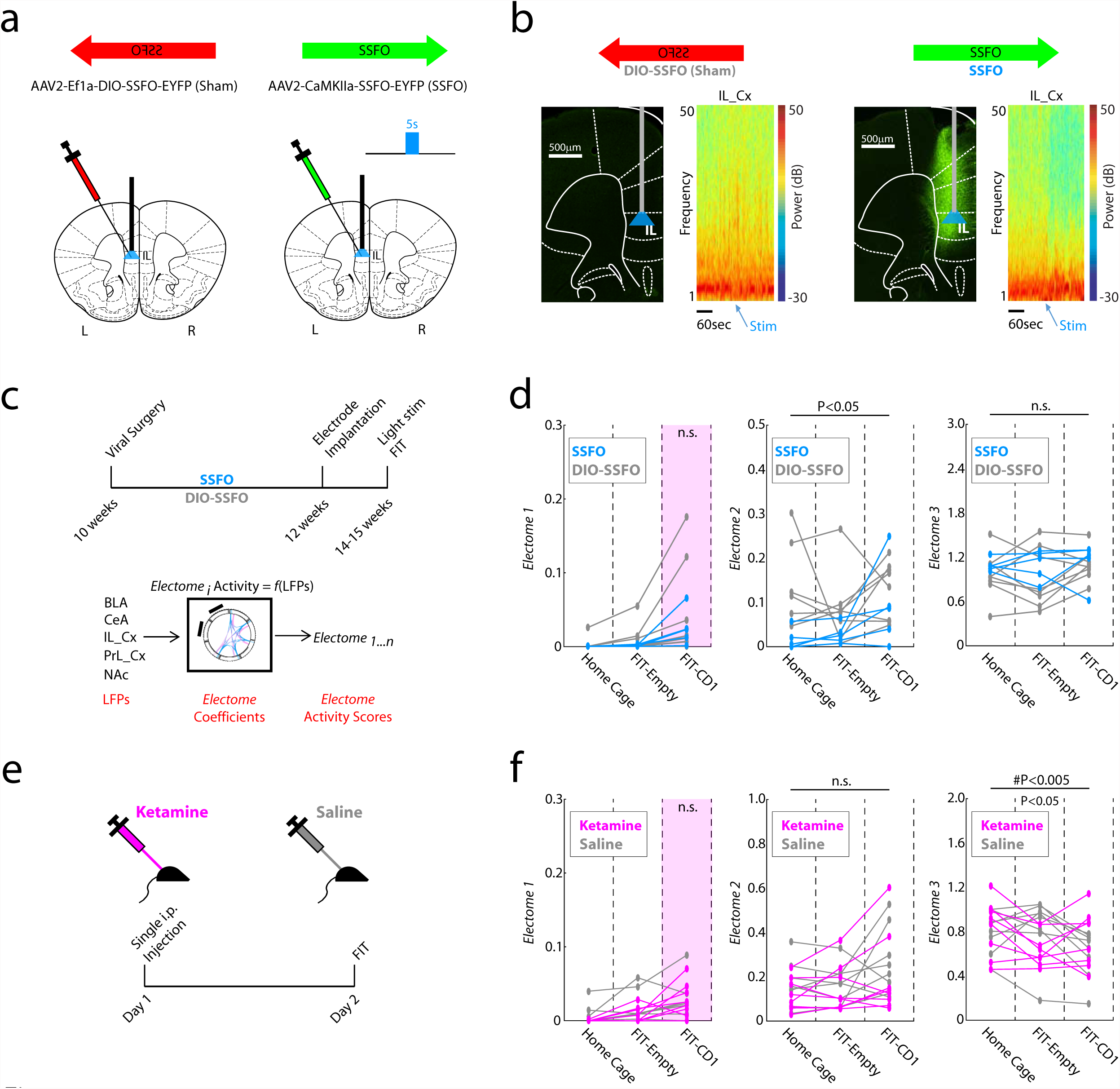
Biologically distinct mechanism underlies depression vulnerability. **a)** IL_Cx infection strategy. **b)** Prominent suppression of IL_Cx gamma (30-50Hz) oscillations was observed after blue light stimulation in animals expressing SSFO. Representative Prefrontal cortex histological images in SSFO mice and DIO-SSFO controls. Broad EYFP labeling was observed in PrL_Cx and IL_Cx in SSFO mice. The light fiber was implanted at the dorsal IL_Cx border. **c)** Schematic for SSFO experiments. **d)** *Electome* activity in SSFO mice compared to the DIO-SSFO sham controls (N=5-8 mice/ group). **e)** Schematic for Ketamine experiment. **f)** Electome activity in Ketamine treated mice compared to saline treated controls (N=8 mice/group).

Ketamine is an emerging rapidly-acting antidepressant agent. A single dose of ketamine (10-20mg/kg, i.p.) has been shown to ameliorate susceptibility in the cSDS model when it is administered after the last defeat episode but 24 hours prior to behavioral testing[33, 34]. Critically, when this same ketamine regimen is given 24-hours prior to the first cSDS defeat episode, there is no effect on an animal’s susceptibility to subsequent cSDS[34]. From a behavioral standpoint, this finding suggests that this ketamine regimen does not target the biological mechanisms underlying vulnerability in C57 mice. Thus, we tested whether ketamine (20mg/kg i.p) was sufficient to suppress *Electome 1* activity, again in stress-naïve mice. Animals were treated with ketamine, and forced interaction testing was performed 24 hours later (Fig. 6e). Applying our initial cSDS Electome model and coefficients to our recorded LFP data, we found that ketamine also failed to suppress *Electome 1* activity (P=0.41 using rank-sum test; N=8 mice/group). However, ketamine did suppress *Electome* 3 activity (F_2,28_=5.61, P=0.005; P<0.05 using post-hoc rank-sum test). *Electome* 2 was not affected by this manipulation (Fig. 6f; F_1,28_=1.27, P= 0.28). Thus, as hypothesized, neither manipulation of post-stress behavioral pathology impacted our stress vulnerability signature, *Electome 1*, providing clear evidence that neural network mechanisms conferring stress vulnerability are distinct from those mediating a stress-induced depression-like behavioral state.

## Discussion

It has been proposed that depression is a brain circuit/network disorder. Using machine learning, we discovered a naturally occurring spatiotemporal dynamic network that signals vulnerability to cSDS in stress-naïve mice (*Electome 1*). We validated this network in three independent models of depression vulnerability (P=6.2x10^−4^ using Fisher’s combined probability test) and demonstrated this network was not affected by our two antidepressant-like manipulations (see supplemental Fig. S5 for summary of findings). Thus, we provide clear evidence that *Electome 1* encodes a convergent depression vulnerability pathway that is biologically distinct from the pathway that signals depression-like behavioral dysfunction. The identification of a convergent depression neural network in stress-naïve mice with normal behavioral function raises the potential that brain spatiotemporal dynamics can be exploited to develop a novel class of therapeutics that actually prevent the emergence of depression in vulnerable individuals in response to stressful experiences.

We discovered two spatiotemporal dynamic networks that together reflect the emergence of depression-like behavior in susceptible animals after cSDS (*Electome 2 and Electome 3*). Two distinct antidepressant-like manipulations each suppressed activity in one, but not both, of these *Electomes. Electome 2 and 3* were only distinguished by their spatiotemporal dynamic features including the frequency of oscillatory synchrony and the directionality of information flow through the regions. Thus, our findings suggest that multiple networks may have to synergize in order to yield a global spatiotemporal dynamic global brain state that mediates depression. Furthermore, suppressing activity in any one of these networks may be sufficient to reverse depression pathology (*i.e.*, an *Electome* two hit model, see supplemental Fig. S6). Alternatively, each of these *Electomes* may mediate a different subset of depression-related behaviors, and suppressing their activity may only normalize these behaviors. Further experiments will be needed to test these hypotheses.

Our three behaviorally relevant and biologically distinct networks (*Electomes 1, 2, and 3*) each involve all of the brain regions we recorded. Future experiments will be needed to clarify the role of additional depression-related brain regions in the *Electome* networks. Nevertheless, since these three *Electomes* were only distinguished by their spatiotemporal dynamic features, our findings establish altered spatiotemporal dynamics across wide-spread neural circuits as a key pathophysiological mechanism underlying emotional disorders.

## Methods Summary

### Animal Care and Use

C57BL/6J (C57) male mice purchased from the Jackson Labs and CD1 male mice purchased from Charles River Laboratory were used for cSDS, Sdk1, and IFNα experiments presented in this study. The C57 mice used for early life stress studies were bred within the Duke Vivarium. C57 mice were housed 3-5 per cage and separated after surgery. All animals were maintained on a 12 hour light/dark cycle, in a humidity- and temperature-controlled room with water and food available *ad libitum*. All CD1 mice were single-housed. Studies were conducted with approved protocols from the Duke University Institutional Animal Care and Use Committee and were in accordance with the NIH guidelines for the Care and Use of Laboratory Animals.

### Neurophysiological Data acquisition

Neuronal activity was sampled at 30kHz using the Cerebus acquisition system (Blackrock Microsystems Inc., UT). Local field potentials (LFPs) were bandpass filtered at 0.5–250Hz and stored at 1000Hz. Neurophysiological recordings were referenced to a ground wire connected to two ground screws.

See supplemental materials for additional methods.

## Acknowledgements

We would like to thank C. Liston and H.S. Mayberg for helpful comments on this manuscript, and Chris Choi for technical support. This work was supported by funding from NIH grant MH79201-03S1 to BDS; NIH grant MH79201 and Lennon Family Foundation to MGC. NIH grant MH099192-05S1 to CB; P50MH096890 to EJN; DARPA HIST program managed by Dr. Jack Judy to LC; One Mind Institute Rising Star Award and R01MH099192 to KDz. We also thank Kerima L. Collier for her generous contribution to KDz. A special thanks to Freeman Hrabowski, Robert and Jane Meyerhoff, and the Meyerhoff Scholarship Program. See supplemental methods for detailed author contributions.

